# Covariate-aware genomic prediction of blood metabolite profiles using multi-task neural networks

**DOI:** 10.64898/2026.06.08.728708

**Authors:** Merve Nur Güler, Estonian Biobank Research Team, Maris Alver, Toomas Haller, Flora Jay, Luca Pagani, Lili Milani, Burak Yelmen

## Abstract

Predictive models of quantitative traits can combine genetic and covariate information, but overall predictive performance alone does not reveal whether differences between models arise from covariate, genetic or joint covariate-genetic effects. Circulating metabolites provide a high-dimensional set of clinically relevant quantitative traits in which these effects can be examined systematically. Although their genetic determinants are well characterised through genome-wide association studies, marginal associations do not establish how accurately metabolomic profiles can be predicted or whether nonlinear models can improve prediction by capturing complex and potentially interactive structure. Here, we developed a multi-task neural network (NN) for simultaneously predicting metabolomic profiles with a three-stage architecture separating covariate, genetic and joint contributions. In comparative analyses, the multi-task NN demonstrated the strongest mean performance across metabolites (R² = 0.219), followed by the single-task NN (R² = 0.211), elastic net (R² = 0.207), and an activation-free multi-task model (R² = 0.191). Decomposition analyses indicated that gains were mainly driven by nonlinear covariate modelling, consistent with improved prediction after incorporating nonlinear age effects into a linear model. Together, these analyses provide a framework for identifying sources of predictive differences between different models, which may be applicable to other collections of quantitative traits.

## Introduction

Circulating metabolites provide proximal molecular readouts of physiological state, reflecting genetic background, environmental exposures and dynamic biological processes^1^. As a vital component of biological systems, the metabolome bridges the genome and the observed phenotype^2^, capturing variation in lipid and amino acid metabolism, inflammation, energy balance and other processes relevant to human health. Metabolites are therefore widely used to characterise cardiometabolic health and stratify disease risk^3^. To achieve stratification at a population level, researchers have turned to biobanks that link expansive metabolite profiles with genome-wide genetic data^4,5^. High-throughput nuclear magnetic resonance (NMR) platforms, including the Nightingale Health platform, have been central to this effort, providing quantitative measurements of lipoprotein lipids, fatty acids, amino acids and other circulating metabolites in a single assay^6^. These datasets not only reveal associations between biomarkers and disease^5–7^ but also allow for the development of genetic predictors^8^, which could be utilised in clinical settings (e.g., HDL/LDL genetic risk estimation), extend metabolite-informed analyses to cohorts without metabolomic assays and support phenome-wide studies of genetically influenced metabolic variation. However, genomics-metabolomics studies have largely focused on mapping the genetic architecture of circulating metabolites, rather than on systematic out-of-sample prediction from genomic data. Genetic studies have identified common-variant associations for conventional lipid traits, including LDL cholesterol, HDL cholesterol, triglycerides and total cholesterol^9^ and metabolomics genome-wide association studies (GWAS) extended this framework beyond broader panels, including lipoprotein subclasses, fatty acids, amino acids and other small molecules^4,10,11^. Biobank-scale studies have further shown that these traits are polygenic, pleiotropic and heterogeneous in SNP heritability^5,12,13^. While these studies have mainly established that circulating metabolites are genetically influenced, it remains unclear how accurately multi-metabolite profiles can be predicted, or whether shared metabolite structure improves prediction over strong single-trait or linear approaches.

In this study, we hypothesised that genomic prediction of metabolites might be improved by multi-task nonlinear modelling for two reasons. First, circulating metabolites are structured rather than independent. Many NMR measures are strongly correlated, particularly within lipoprotein subclasses and related lipid measures, because they describe related properties of the same particles and broader metabolic states^6^. Extensive pleiotropy^14^ and shared physiological exposures may further reinforce this covariance, allowing the metabolome to be described by a limited number of underlying biological dimensions^6^. Multi-task learning^15^ may therefore improve efficiency by allowing related phenotypes to share signal. Although multi-task deep learning for metabolites is scarce, evidence from other domains support this premise: multi-task learning can improve parallel estimation of polygenic predictors when targets share genetic basis^16^ and improve multi-trait genomics prediction involving non-additive interactions^17^.

Second, part of the unexplained phenotypic variation might be explained by interaction effects including epistasis^18^ between genetic variants, modulation of variant effects by the broader polygenic background^19^, or interactions between variants and covariates such as age, sex and body mass index (BMI)^20,21^. Furthermore, covariates may themselves act nonlinearly and interactively. Previous work has demonstrated that modelling nonlinear and interactive covariate effects can improve phenotypic prediction and association power without sacrificing performance when effects are effectively linear^22^. While nonlinear neural network (NN) architectures possess the theoretical capacity to capture these high order interactions without explicit prior specifications^23^, translating this into superior predictive performance has proven challenging in practice. Genomic prediction benchmarks emphasise that nonlinear models do not automatically outperform strong linear baselines^24,25^. This discrepancy underscores two important points we must remain cautious of. First, finding the optimal model for a specific task is not trivial due to the infinitely large architecture and hyperparameter search space, demanding exhaustive and often domain-specific optimisation^26,27^. Second, sample size acts as a strict constraint; without large-enough datasets, NNs might fail to detect nonlinear effects, and instead overfit to high-dimensional noise rather than capturing true biology^23,28,29^.

Accordingly, we developed an NN framework which (i) separates covariate, genetic and joint covariate-genotype components of metabolite prediction, (ii) exploits cross-metabolite sharing in a multi-task setting, and (iii) benchmarks against strong linear baselines to assess overall and component-specific performance gains. Our findings suggest that multi-task NNs can improve metabolite prediction over state-of-the-art linear baselines, which tends to mainly stem from the nonlinear modelling of covariate features and potentially from shared structure between metabolites. Although evaluated here using metabolomic traits, this framework is not metabolite-specific and could be applicable to other phenotype collections for distinguishing sources of predictive improvement.

## Results

Using genotype and NMR metabolomics data from Estonian Biobank^30^ (EstBB) participants, we evaluated the multi-task prediction of circulating metabolite profiles. After filtering steps (see **Methods**), individuals were split into training, validation, model-selection and held-out test sets, with the held-out test set reserved exclusively for final evaluation. Metabolite-specific GWAS performed in the training set across the full 249-biomarker NMR panel were used to construct an LD-pruned union set of genetic predictors and assess their sharing across metabolic traits. Prediction analyses used genotype data together with age, sex, BMI and the first ten genetic principal components as covariates. They were restricted to 109 traits: 107 non-derived metabolites together with high-density lipoprotein cholesterol (HDL_C) and low-density lipoprotein cholesterol (LDL_C), which were retained because of their clinical relevance. We then compared linear baselines, single-task and multi-task NNs using the held-out test set for final performance evaluation (see **Figure 1**, **Methods**). Linear baselines include the activation-free multi-task NN which retained the same shared multi-output architecture as the nonlinear multi-task NN but removed nonlinear activation functions, serving as an architecture-matched comparator for nonlinearity.

**Figure 1:**
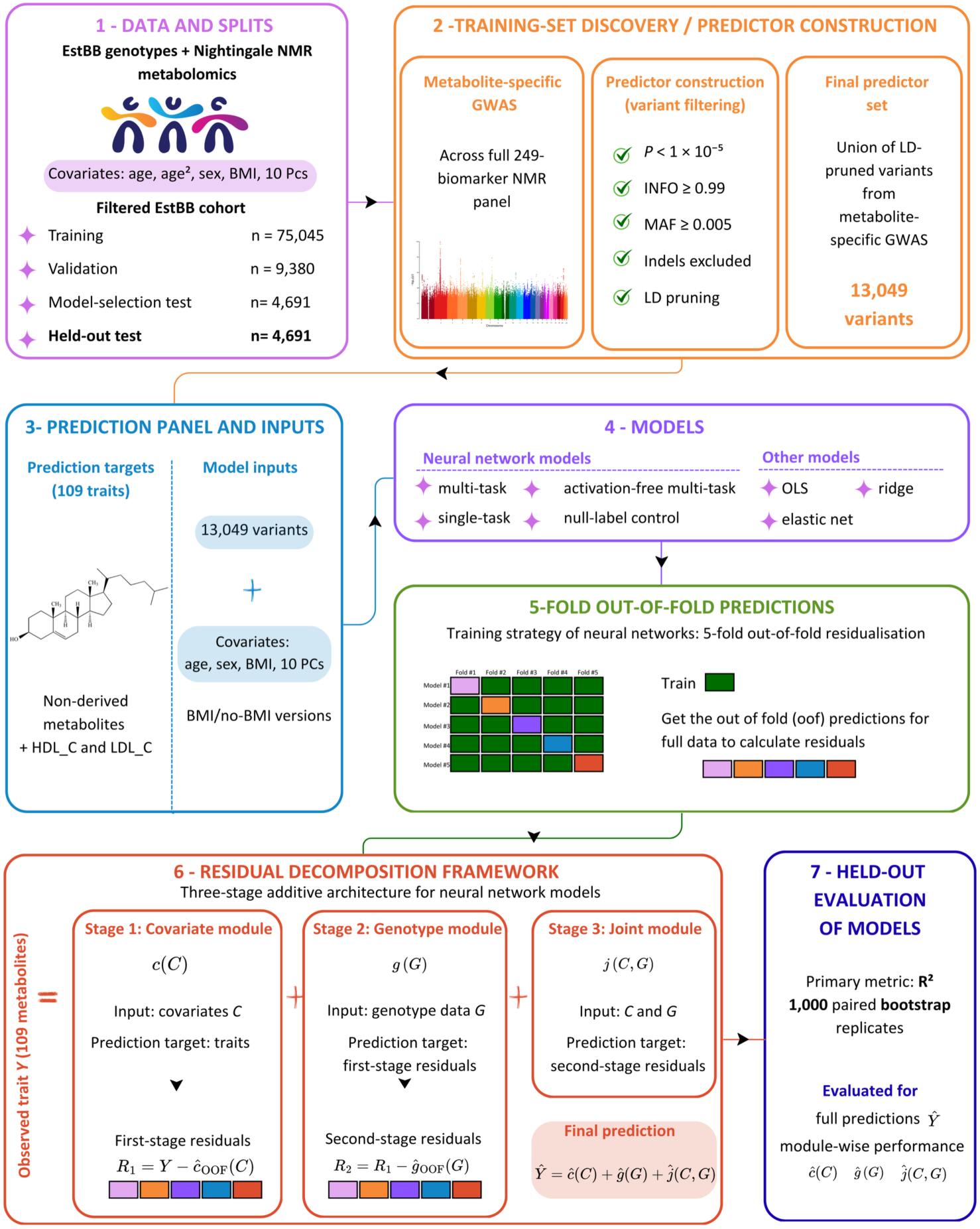
Overview of the study design and modelling framework. Genotype and Nightingale NMR metabolomics data from the Estonian Biobank were split into training, validation, model-selection test and held-out test sets. Metabolite-specific GWAS were performed in the training set across the full 249-biomarker NMR panel, and variants passing the filtering criteria were LD-pruned and merged into a union predictor set of 13,049 variants. Prediction analyses were then restricted to 109 target traits, consisting of non-derived metabolites together with HDL-C and LDL-C. Models used the genetic predictor set together with covariates, including age, sex, BMI and the first ten genetic principal components, with additional BMI-excluded (no-BMI) version. Neural-network models were fitted using a three-stage residual decomposition framework: first a covariate module, then a genotype module trained on first-stage residuals, and finally a joint covariate-genotype module trained on second-stage residuals. Five-fold out-of-fold predictions within the training set were used to construct residual targets and reduce within-sample bias. Final model performance was evaluated only in the held-out test set using full predictions and component-wise predictions, with R² as the primary metric and uncertainty estimated by 1,000 paired bootstrap replicates.

### Cross-metabolite sharing of the genetic predictor set

We defined the genetic predictor set as the union of genetic variants associated with at least one biomarker at P<1×10^-5^ in training-set GWAS across the full 249-biomarker panel. After filtering and LD-pruning, this yielded 13,049 variants (see **Methods**). Using the same association threshold, we characterised cross-metabolite sharing by counting how many biomarkers each LD-pruned variant was associated with. Sharing was substantial: the median number of associated biomarkers per variant was three, with a strongly skewed distribution (**Figure S1a**). Overall, 63.1% of the variants were associated with >1 trait and the variants in the top 1% of the distribution were associated with >157 biomarkers.

### Comparison of overall performance across models

We first compared model performance across 109 prediction traits in the held-out test set (**Figure 2a-2b, Table S1**). The multi-task NN achieved the highest overall performance, with a mean held-out R² of 0.219 across metabolites, compared with 0.211 for single-task NN, 0.207 for elastic net, 0.191 for the activation-free multi-task model, 0.172 for ridge regression, and 0.076 for ordinary least squares regression (OLS). The null multi-task control, trained on permuted labels to provide a baseline, yielded near-zero performance as expected (mean R² = −0.001). Uncertainty in mean performance was estimated using paired bootstrap resampling of held-out test individuals (see **Methods**), where each bootstrap replicate used the same resampled individuals across all models so that model predictions remained paired within individuals. Mean performance and 95% confidence intervals are shown in **Figure 2b**.

**Figure 2:**
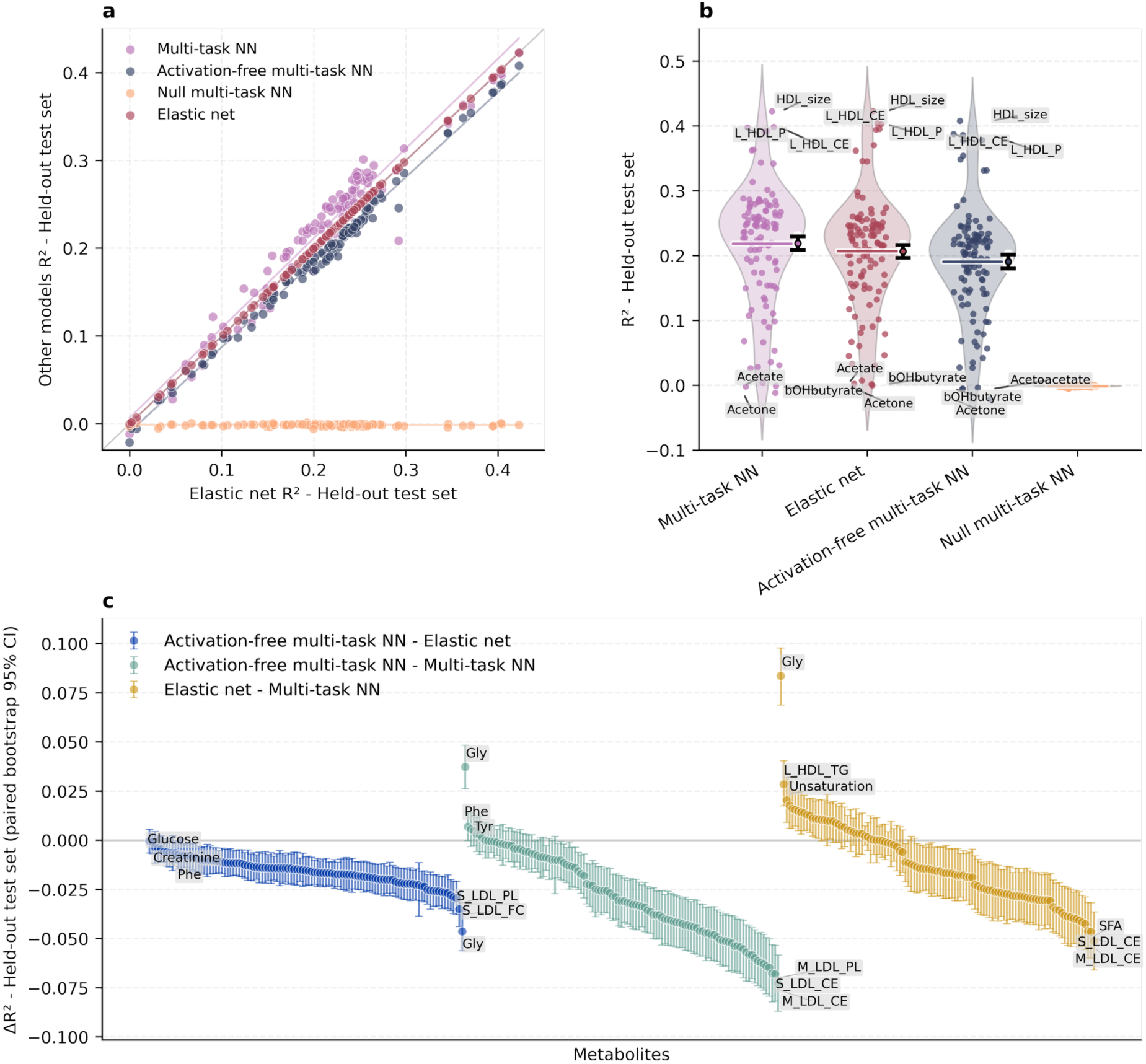
Comparative performance of linear and neural network models across metabolites. The held-out test set performance based on R^2^ is compared over 109 metabolites for different models: multi-task neural network (NN), activation-free multi-task NN, elastic net, and null multi-task NN trained on permuted labels. **a)** The metabolite-level comparison of held-out test-set R² between elastic net (x-axis) and the multi-task models (y-axis), including the null multi-task control. Each point represents one metabolite. The diagonal line indicates equal performance to elastic net. The overlaid lines represent least-squares regression fits across metabolites and are included as visual guides to the average model-specific relationship. **b)** Distribution of the held-out test-set R² values (y-axis) across metabolites for the same models (x-axis), shown as violin plots with individual metabolite values overlaid. Horizontal bars indicate model means. Error bars on mean values represent 95% confidence intervals estimated using paired bootstrap resampling of individuals in the held-out test set. Metabolite names are shown for extreme values to identify traits with particularly high or low prediction performance within each model. **c)** Metabolite-level (x-axis) differences in held-out test-set R² values (y-axis) for selected model pairs. Points represent mean differences estimated using paired bootstrap resampling of individuals in the held-out test set. Error bars indicate 95% confidence intervals derived from the bootstrap distribution. Positive values indicate higher performance of the first model in each comparison. Within each comparison, metabolites are ordered by mean ΔR², so the x-axis is used only for visual separation and has no biological ordering. Metabolite names are shown for the 3 largest positive and 3 largest negative differences within each comparison.

At the metabolite level, paired bootstrap comparisons showed that the multi-task NN outperformed elastic net, the best-performing linear model, for 57 of 109 metabolites (52.3%) based on R², whereas elastic net performed better for 9 (8.3%), and no clear difference was observed for 43 metabolites (39.4%) (**Figure 2c, Table S2-3**). Within the NN models, the nonlinear multi-task NN outperformed the activation-free multi-task model for 85 metabolites (78.0%), with the reverse observed for 2 metabolites (1.8%) and no clear difference for 22 (20.2%). The comparison with the single-task NN was comparatively less decisive: the multi-task model outperformed for 44 metabolites (40.4%), the single-task NN for 4 (3.7%) and 61 metabolites (56.0%) showed no clear winner. These comparative patterns were similar for Pearson’s r, MSE and MAE, indicating that the performance hierarchy was not metric-specific (**Table S1, S2**).

Because BMI can improve prediction but may also lie on genotype-metabolite pathways^31,32^, we repeated the analyses with and without BMI (**Figure S2**, **Table S3**). Since comparative results were similar, with higher predictive power for all models using BMI, we retained BMI in covariates for the main analyses, consistent with a predictive rather than causal modelling objective^33^.

We next examined the difference between the training and the held-out test performance as a descriptive assessment of generalisability. Models trained on the true phenotype labels generally performed better on the training set than on the held-out test set as expected, consistent with some degree of overfitting. This gap was largest for OLS, which also showed poor held-out performance, whereas regularised linear models and NN models showed smaller discrepancies. Overall, these results indicate that model comparisons on the held-out test set were not dominated by an obvious overfitting pattern in the best-performing models. (**Figure S3**).

### Module-wise contributions to held-out predictive performance

To better understand where the predictive gains originated, we decomposed the held-out test-set performance into covariate, genetic, and covariate-genetic joint modules across NN architectures (**Figure 3, Table S3-4**). In this decomposition, the covariate module used age, sex, BMI and genetic PCs, the genetic module used the LD-pruned variant set, and the joint module used both covariates and genetic variants to model remaining residual structure. Metabolite-level architecture comparisons were assessed using paired bootstrap resampling of held-out test individuals (see **Methods**, **Figure 1**).

**Figure 3.**
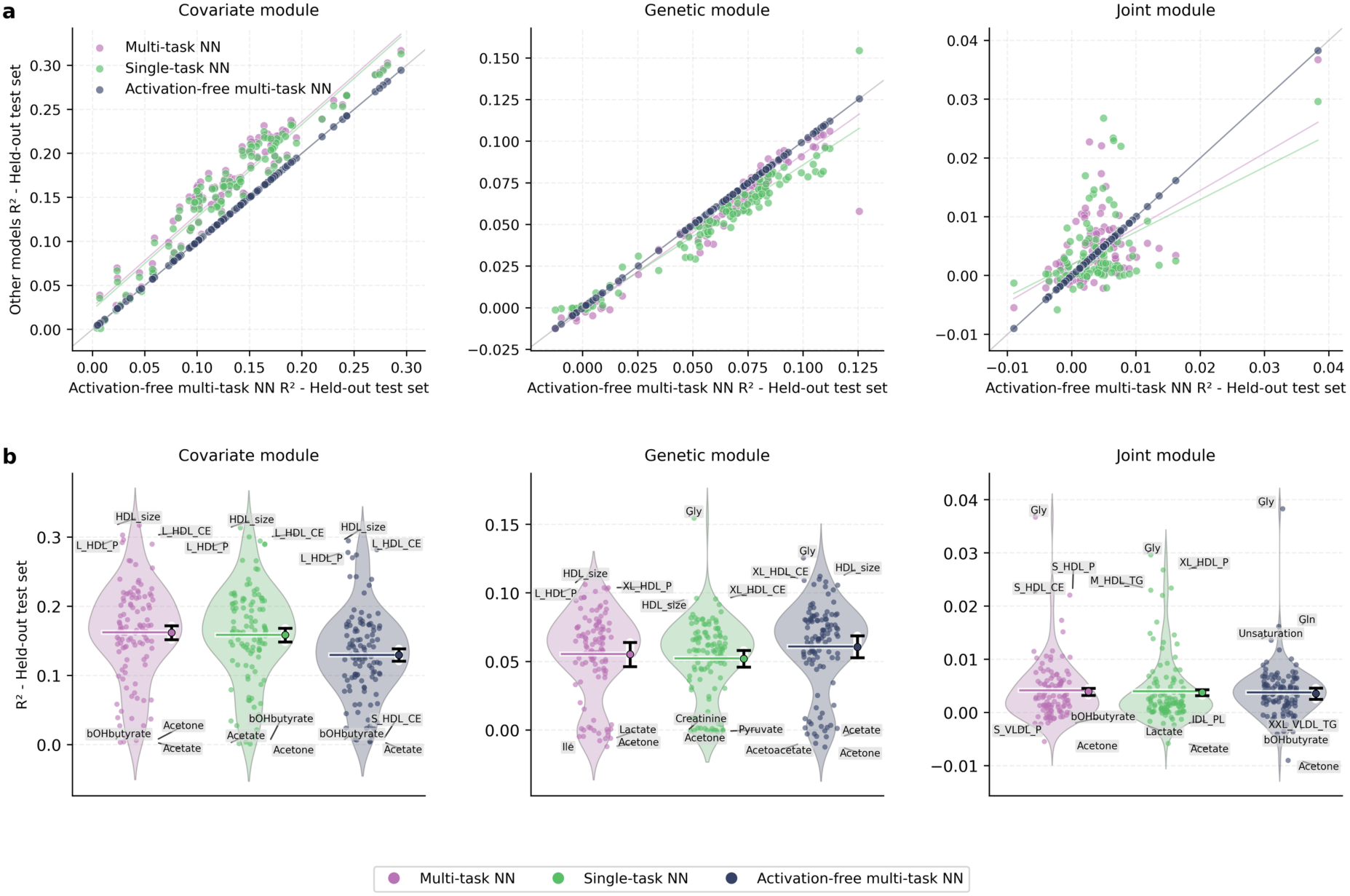
Residual decomposition of held-out prediction across neural network architectures. **a)** Metabolite-level comparison of component-specific held-out test-set R² between the activation-free multi-task neural network (NN) (x-axis) and the other NN architectures (y-axis). Separate panels show covariate (left column), genetic (middle column) and joint (right column) modules. Each point represents one metabolite, and lines indicate fitted linear trends. The diagonal indicates performance equal to activation-free multi-task NN. **b)** Distribution of component-specific held-out R^2^ values (y-axis) across metabolites for the same three architectures (x-axis), shown separately for the covariate (left column), genetic (middle column) and joint (right column) modules. Violin plots show the distribution across metabolites, with individual metabolite values overlaid. Horizontal bars indicate model means and error bars indicate bootstrap-derived 95% confidence intervals for the mean across metabolites. Metabolite names are shown for extreme values to identify traits with particularly high or low prediction performance within each model. Component-specific R² values were calculated against the observed phenotype rather than against the residual target used to train each module; thus, they quantify how strongly each component prediction aligns with phenotype variation on the original prediction scale.

For this module-wise analysis, we used the activation-free multi-task NN as the linear comparator rather than elastic net. Elastic net was the strongest conventional linear baseline overall but applying it in this residual-decomposition framework would have required repeated cross-validated out-of-fold fitting at each stage and for each metabolite, making it computationally prohibitive (**Figure S4**). The activation-free multi-task NN retained the same decomposition procedure, and the same inputs and output structure as the multi-task NN, but removed nonlinear activation functions, thereby providing a direct comparator within the same framework.

The covariate module captured the largest component-level signal across all architectures. The mean held-out R² for the covariate module was 0.162 for the multi-task NN, 0.159 for the single-task NN, and 0.130 for the activation-free multi-task NN (**Figure 3, Table S4**). In direct metabolite-level comparison, the multi-task NN had higher covariate-module R² than the activation-free multi-task NN for 105 of 109 (96.3%) metabolites, and paired bootstrap analysis identified a clear advantage for 100 of 109 (91.7%) metabolites (**Figure 4, Table S5**). The comparison between the multi-task and single-task NNs was closer: the multi-task NN was favoured for 48 metabolites (44%), while the remaining showed no clear winner.

**Figure 4.**
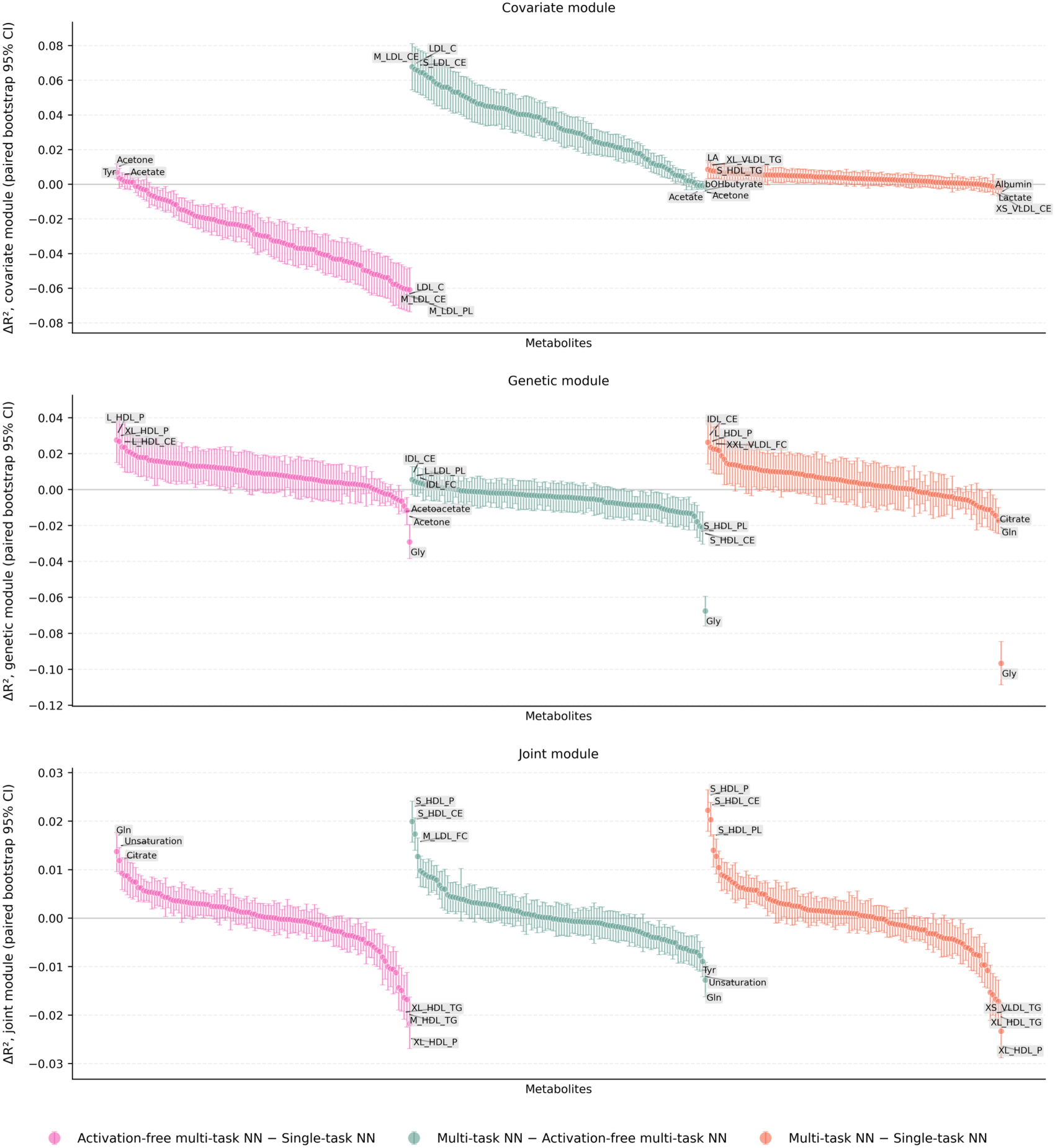
Paired bootstrap comparison of component-specific performance differences between neural network architectures. For each decomposition component, points show the metabolite-specific mean difference in held-out test-set R² (y-axis) between model pairs (x-axis) and error bars indicate 95% confidence intervals from paired bootstrap resampling of individuals. Panels show differences for the covariate module, genetic module and joint module, respectively. Component-specific R² values were calculated against the observed phenotype rather than against the residual target used to train each module; thus, they quantify how strongly each component prediction aligns with phenotype variation on the original prediction scale. Positive values indicate better performance of the first model named in each comparison. Within each panel, metabolites are grouped by model comparison and ordered by mean ΔR²; therefore, the x-axis position is used only for visual separation and has no biological ordering. Metabolite names are shown for the 3 largest positive and 3 largest negative differences.

The genetic module contributed a smaller but measurable increase in explained variance across all models. The mean module-alone genetic R² was 0.055 for the multi-task NN, 0.061 for the activation-free multi-task NN and 0.052 for the single-task NN (**Figure 3**, **Table S4**). The multi-task NN was favoured over the single-task NN for 19 metabolites (17.4%), the single-task NN for 11 metabolites (10.1%), and 79 metabolites (72.5%) showed no clear winner. Interestingly, the activation-free multi-task NN outperformed the nonlinear multi-task NN for 32 metabolites (29.4%) but ΔR² values were small, and there was no clear winner for 77 metabolites (70.6%) (**Figure 4, Table S5**).

The joint module showed the smallest and least stable contribution across all models. The mean module-alone joint R² was 0.009 for the multi-task NN, 0.004 for the activation-free multi-task NN, and 0.004 in the single-task NN (**Figure 3**, **Table S4**). Incremental gains from this stage were also small: the mean ΔR² from adding the joint module was 0.003 for the multi-task NN and close to zero or slightly negative for the activation-free and single-task models (**Table S4**). Paired bootstrap comparisons did not show a consistent architecture-specific advantage. For the joint module, the multi-task NN was favoured over the activation-free model for 21 metabolites (19.3%), the activation-free model for 23 metabolites (21.1%) and 65 metabolites (59.6%) showed no clear winner. Similarly, the multi-task NN was favoured over the single-task NN for 27 metabolites (24.8%), the single-task NN for 25 metabolites (22.9%) and 57 metabolites (52.3%) showed no clear winner (**Figure 4, Table S5).**

As an additional check on covariate-associated nonlinearity, we fitted a linear model using only age and compared it to a linear model fitted using age and age². The quadratic term improved performance across most metabolites (mean ΔR² ≈ 0.013; 98 of 109 metabolites (90%)). This supports the interpretation that nonlinear extensions can capture additional covariate-related variation and likely contribute to the advantage of the multi-task NN (**Figure S5**).

Overall, the module-wise analysis showed that the nonlinear multi-task NN achieved the highest full-model performance (mean R² = 0.219), despite not being the top-performing architecture consistently for every individual module. Its clearest advantage was observed in the covariate module relative to the activation-free comparator, while genetic module effects were present but heterogeneous over metabolites. The joint module contribution was similarly heterogeneous, but its contribution to the prediction was limited over all models.

### Metabolite-level genetic and joint module gains

We then examined whether component-wise gains were driven by only a small number of extreme metabolites or reflected a broader pattern across the panel (**Figure S6, Table S3**). The addition of the genetic module increased the held-out R² for most metabolites in all three architectures: 97 of 109 metabolites (89.0%) in the multi-task NN, 102 of 109 (93.6%) in the activation-free multi-task NN and 99 of 109 (90.8%) in the single-task NN. In contrast, gains from the joint module were smaller and less consistent. The median joint-module gain was close to zero in all three models, indicating that this stage did not provide a broad additional improvement across the panel.

### Observed genetic prediction performance relative to SNP-heritability-based expectations

To assess how closely models approach the approximate theoretical limit imposed by genetic architecture (see **Methods**), we compared observed genotype-based correlations with expectations derived from SNP-heritability (**Figure S7a, Table S6**). Across metabolites, observed correlations were generally below this bound, indicating incomplete recovery of the additive genetic signal. The gap was prevalent across models, which suggests that the limitation might not only be model capacity but potentially also finite sample size, imperfect tagging of causal variants due to the reduced SNP set, and residual phenotype noise. We also compared the same expectations with full-model predictive correlations (**Figure S7b**), which demonstrated that even for low-heritability metabolites, theoretical predictive limit can be surpassed with the addition of covariates.

## Discussion

Circulating metabolites measured by NMR provide a clinically relevant and system-level assessment of human physiology, capturing both genetic and environmental influences. These panels have been widely used for disease risk prediction and epidemiological profiling yet predictive modelling has largely treated metabolites as independent outcomes. To address this gap and test whether shared modelling can improve prediction, we modelled metabolites jointly and evaluated whether multi-task NNs improve genome-based prediction across a large circulating metabolite panel relative to linear baselines. We further decomposed the predictive models into covariate, genetic and combined covariate-genotype signals to pinpoint the source of potential gains. Large-scale metabolite GWAS have shown that circulating metabolic traits have measurable but heterogeneous SNP-heritability, indicating that part of inter-individual metabolic variation is genetically anchored. Consistent with this, the genetic predictor set used here showed dense sharing across metabolites (**Figure S1**), providing an empirical basis for a multi-task NN that can use shared hidden layers to model cross-metabolite structure.

It is important to first underline the differences in our comparison framework to place our findings in appropriate context. Relative to elastic net (and other simpler linear baselines, OLS and ridge regression), the nonlinear multi-task NN represents the complete modelling package: joint prediction of metabolites, shared hidden layers across outputs, and nonlinear transformations. Relative to the activation-free multi-task NN, it tests the added value of nonlinear transformations within the same multi-output framework. Relative to the single-task NN, it speaks to the value of modelling metabolites jointly, although this comparison is less concrete because the single-task models were not optimised to the same extent. In this context, the nonlinear multi-task NN achieved the strongest held-out performance overall compared to other models, but the improvement over both single-task NN and elastic net was modest. In bootstrap comparisons, the multi-task NN outperformed elastic net for 57 of 109 metabolites (52.3%), while elastic net performed better for 9 (8.3%) and 43 (39.4%) showed no clear winner. In addition, it had superior performance over the activation-free multi-task model, with better performance for 85 metabolites (78%). By contrast, the comparison with the single-task NN was closer: the multi-task NN was favoured for 44 metabolites (40.4%), the single-task NN for 4 (3.7%) and rest of the metabolites showed no clear winner. Thus, the conclusion is not that multi-task NN is uniformly superior, but that it gives the strongest average held-out performance while preserving substantial metabolite-level heterogeneity.

Decomposing metabolite prediction into covariate, genetic and joint covariate-genotype components provides a more informative view of where this advantage comes from. Module-wise analyses indicated that the clearest architecture-dependent difference occurred in the covariate module. The nonlinear multi-task NN and single-task NN outperformed the activation-free multi-task NN for the covariate component across nearly all metabolites, suggesting that nonlinear modelling of covariate-associated structure contributed substantially to the full-model advantage. This is consistent with previous work showing that flexible modelling of nonlinear and interactive covariate effects can improve phenotype prediction and association analyses^22^. In our setting, this likely reflects nonlinear age trajectories, sex-specific effects, BMI-related structure and higher-order covariate dependencies that are not captured by simple linear covariate terms^22,34–36^. The improvement observed after adding age² to linear models supports this interpretation, showing that even simple nonlinear extensions can provide additional signal (**Figure S5**).

By contrast, the genetic module predictions were broadly similar between the nonlinear multi-task and single-task models, and the activation-free multi-task model had slightly higher average genotype-component R². The joint module, designed to capture remaining structure after covariate and genotype main effects, contributed little across all architectures. Taken together, these results do not support a dominant role for higher-order genetic or genotype-covariate interaction effects in driving the observed performance gains, at least within the present sample size, feature set and optimisation regime. However, there was some variability across the panel, indicating that these effects might be more relevant for some metabolites (**Figure 4, Figure S6**).

The SNP-heritability-based comparison provides a separate perspective on the genotype-derived component. The observed genotype-derived correlations remained below the approximate heritability-implied expectation for most metabolites across both linear and nonlinear NN models, indicating that a substantial fraction of additive genetic signal remains unrecovered (**Figure S7**), but this gap should not be interpreted as a failure of the models alone. The reference is based on genome-wide SNP-heritability estimates, whereas our prediction models used a GWAS-selected and LD-pruned subset of variants. Finite sample size, imperfect tagging of causal variants, and noise could all reduce the recoverable signal. The heritability comparison therefore places the results in context: multi-task learning might improve predictive efficiency, but it does not remove the central constraints of genomic prediction.

The biological grouping of metabolites showed that model performance was strongly structured across the metabolome, although these comparisons should be interpreted descriptively because group sizes were highly uneven, especially for lipoprotein subclasses. With that caveat, the clearest gains of the nonlinear multi-task NN were concentrated in lipid and lipoprotein-related traits, particularly lipoprotein subclass measures, cholesterol traits, fatty acids and other lipids (**Figure S8, Table S3)**. Among the 25 metabolites with the largest multi-task NN gains over elastic net, 17 were lipoprotein subclasses, 3 were fatty acids, 2 were cholesterol-related measures, 2 were other lipids and 1 was an apolipoprotein. Importantly, this lipid-centred pattern was also visible when the multi-task NN was compared with the single-task NN: the largest multi-task gains were again concentrated in lipoprotein subclasses and other lipid-related measures. This pattern is biologically coherent. Many lipid measures in the NMR panel describe related properties of lipoprotein particles, so their covariance reflects shared particle composition and lipid-transport biology rather than independent metabolite-specific signals^6,37,38^ which might explain the advantage of joint modelling. The group-wise comparisons also suggest that the observed advantage cannot be attributed to multi-task learning alone. The activation-free multi-task model was generally less competitive with elastic net, whereas the nonlinear multi-task and single-task models showed clearer gains, particularly for lipid and lipoprotein traits. Thus, the lipid-specific improvement potentially reflects a combination of shared metabolite structure and nonlinear modelling.

Several design choices influence how our findings should be interpreted. First, the predictor set was restricted to GWAS-selected and LD-pruned variants to reduce the dimensionality of genomic input^39^. This made the NN training tractable, but it also means that the models operated on a curated subset of genetic signal rather than the full genome and may miss weak, distributed effects. Because genetic predictors were selected using marginal additive GWAS associations, variants whose effects are mainly interaction-driven or weakly distributed may be underrepresented. This feature-selection step may favour additive genetic signal and partly explain why the activation-free model was competitive in the genetic module. Second, the models were trained and evaluated in unrelated European-ancestry individuals from EstBB. Although this reduces population-structure heterogeneity, it limits ancestry generalisability. External validation in more diverse cohorts is therefore required. Third, covariate inclusion, particularly BMI, introduces a trade-off between prediction and interpretation. BMI is a useful predictor of circulating metabolites, but it is not a neutral adjustment variable: it is itself partly genetically determined and closely linked to broad metabolic differences associated with adiposity and obesity^31,32^. Conditioning on BMI may therefore improve predictive performance while also changing the apparent contribution of genetic effects to metabolite variation. However, despite that the biological interpretation remains nuanced, the stability of model rankings with and without BMI suggests that the main predictive conclusions are robust (**Figure S2**). Fourth, additional physiological states and environmental exposures were not explicitly modelled, which may add noise and mask subtle genetic or interaction effects. However, if the main goal is predictive power, the covariate module can be extended, or additional modules can be integrated to the decomposition framework. Fifth, hyperparameter optimisation differed between multi-task and single-task models because exhaustive metabolite-specific tuning was computationally impractical, which reflects a real constraint in genomic deep learning and limits the strength of direct architecture comparisons. Finally, activation-free NNs are not identical to conventional linear regression; optimisation choices, regularisation and batch normalisation can still affect their behaviour^40^, which affects the interpretation of our module-specific findings. We used activation-free NN, the second-best linear model, as a proxy for linear modelling because decomposition was not computationally feasible for elastic net (**Figure S4**), which was the best-performing linear model (**Figure 2**). For this reason, the observed module-specific performance advantages of nonlinear multi-task model cannot be assigned cleanly to multi-task learning alone or to nonlinearity alone.

More broadly, these results highlight a practical point that is often underemphasised: optimisation itself is a major bottleneck in deep learning for genomics. Multi-task learning reduces this burden by allowing a single model to be tuned once and applied across many traits, whereas single-target approaches require repeated optimisation. In this sense, the advantage of multi-task models is both statistical and computational. At the same time, the modest improvement over elastic net is a useful reminder that strong linear baselines remain difficult to beat in genomic prediction^23–25^. Neural networks should therefore be evaluated not only by whether they improve average prediction, but also by which factors they improve, which signals they use and whether the additional complexity changes the biological interpretation.

In this regard, interpretability is an important direction for future work. Although the modular decomposition used here showed that most gains arose from covariate-associated and shared lipid structure, it does not identify the specific variants, loci, pathways or covariate patterns driving metabolite-specific predictions. Post hoc attribution methods could therefore be applied to the genetic variants and covariate inputs to interpret the features used by the model, further pinpointing the advantages/disadvantages of neural networks and providing additional biological context^41^. A complementary approach would be to incorporate biological structure directly into the architecture, for example through gene-, pathway-or ontology-informed connectivity^42^, as in biologically informed neural-network models^43,44^. The present model was instead kept as a flexible data-driven architecture, without pathway or lipoprotein-hierarchy constraints. This preserves the model power but limits mechanistic interpretation. Future hybrid approaches could combine the flexibility of multi-task prediction with more interpretable biological organisation, although such constraints should be used carefully because they may also over-encode domain knowledge and reduce opportunities for novel discovery.

A natural translational direction is to treat these models as metabolite-level genetic score generators. Predicted metabolite profiles are not substitutes for measured metabolomics, especially for state-sensitive metabolites, but they could extend metabolite-informed analyses to cohorts without NMR profiling and support phenome-wide studies of genetically influenced metabolic variation. This may be most realistic for the structured lipid and lipoprotein traits where prediction was strongest, including HDL-, LDL-and VLDL-related measures. In this context, the advantage of multi-task NN models would not only be potentially improved predictive power but also the ability to predict complete NMR metabolome profile with a single compact model. Since inference after training can be done in seconds, genetic scores for the complete NMR metabolome profile, or a set of disease-relevant metabolites, could be obtained for better individualised medical interventions. However, before such models can be used for these purposes, they will require external validation, calibration across cohorts and explicit testing of downstream utility. External validation should test transfer across cohorts, ancestries and metabolomics platforms. UK Biobank provides a natural first validation setting because genotype data and Nightingale NMR metabolomic measurements are available in the same large population cohort^6^. Validation in UK Biobank or other European-ancestry biobanks using the same NMR platform would test cohort transfer under comparable assay conditions, whereas cohorts with different ancestry composition or alternative metabolomics assays would be needed to quantify population and platform shift.

In summary, nonlinear multi-task NNs provided the strongest average held-out performance for genome-based prediction of circulating metabolites, but the improvement over single-task NNs and elastic net was modest and metabolite-dependent. This gain can be primarily attributed to improved modelling of covariate-associated and shared structure between metabolites, whereas contributions from genotype-only and joint genotype-covariate structure were small, heterogeneous across metabolites and did not outperform linear modelling consistently. Larger sample sizes, richer environmental measurements and more interpretable architectures could help further define the extent of this observed advantage and how well it translates into improved downstream biological and clinical applications.

## Methods

### Study population and data

Genotype data (n = 211,259) were obtained from EstBB^30^ GSA-array framework (Illumina’s GSA-MD-24v1, GSA-MD-24v2, ESTchip1_GSA-MD-24v2, ESTchip2_GSA-MD-24v3, 2023-0612 snapshot). We used the non-imputed genotype data for linkage disequilibrium (LD) calculation and for the first step of GWAS. We used the imputed genotype data, generated via Estonian reference panels and the Haplotype Reference Consortium (HRC) panel, for GWAS and for selecting the final SNP set for prediction models. In the analysis, we included age, sex, body mass index (BMI) and the first ten genetic principal components (PCs) as covariates. Blood circulating metabolite levels were obtained from nuclear magnetic resonance (NMR) metabolomics measurements generated using the Nightingale platform from EDTA plasma samples. The Nightingale platform provides quantitative measurements for a panel of metabolic biomarkers, including lipoprotein lipids, fatty acids, amino acids and other small molecules^14,37^. In total, 249 metabolite biomarkers were available. Of these, 107 were non-derived biomarkers representing directly measured metabolite concentrations, while the remaining 142 were derived biomarkers calculated as ratios or composite measures of the primary metabolites.

### Filtering and dataset splitting

We first restricted the analysis to a genetically homogeneous population. We excluded individuals with non-European ancestry based on ancestry grouping estimated with *bigsnpr*^45,46^ using imputed genotype data. We then removed individuals related up to the second-degree using relatedness estimates from *KING*^47^ v2.2.7. We also excluded individuals without available NMR metabolomics measurements. When multiple NMR measurements were available for an individual, we selected the sample with the fewest missing metabolite values. For individuals with multiple BMI records (GEVA snapshot 2025v01), the BMI measurement closest in time to the metabolite sampling date was selected. After these filtering steps, we randomly split the remaining individuals into four datasets: training (n =75,045), validation (n =9,380), model-selection test (n =4,691) and held-out test (n =4,691). We used the validation and model-selection test sets while developing and selecting the models, and we kept the held-out test set only for the final evaluation of model performance.

### Calculation of genetic principal components

Population structure was accounted for by including genetic principal components (PCs) as covariates in the analyses. To avoid leakage, PCs were computed using only the training dataset with *PLINK2*^48^ *v2.00a3LM*. Principal component analysis (PCA) was performed on genotype data using variants with minor allele frequency (MAF) ≥0.01. Prior to PCA, variants were pruned for LD using a sliding window approach (window size 100 variants, step size 10 variants, r^2^ <0.1). SNP loadings obtained from the training PCA were then used to project individuals from the validation and test datasets onto the training-derived PC axes, ensuring that all samples were represented in the same PC space. Because PC scores obtained through projection are known to be biased toward zero in high-dimensional settings, a shrinkage correction was applied following the method described by Lee et al^49^, which rescales projected PC scores based on the ratio of variants to samples used in PCA. The first ten PCs were included as covariates in all downstream analyses.

### GWAS for variant selection

We performed GWAS to identify variants associated with each metabolite level only with the training set, to prevent information leakage between dataset splits. Association testing was carried out using *Regenie*^50^ *v3.4* within its two-step ridge regression framework. In step 1, we fitted a whole-genome regression model using array genotype data. Before model fitting, we filtered variants with *PLINK2 v2.00a3LM* to retain markers with MAF≥0.01, minor allele count ≥20, Hardy– Weinberg equilibrium P >1×10^−15^, genotype missingness ≤0.1, and individual missingness ≤0.1. This step captures the background polygenic signal and produces leave-one-chromosome-out (LOCO) predictions for each phenotype. In step 2, we tested single-variants using imputed genotype data. We retained imputed variants with INFO score >0.99 and MAF ≥0.005. Association statistics were calculated under an additive genetic model for each metabolite while conditioning on the LOCO predictions from step 1. Covariates included age, age-squared, sex, and genetic PCs. *Regenie* handled missing genotype, phenotype and covariate values using its default settings (mean-imputation).

### Construction of the genetic predictor set for prediction models

After GWAS, we extracted variants from the *Regenie* step 2 results for each metabolite at several significance thresholds (−log_10_((P) >4, 5, 6, and 7, as well as the genome-wide significance threshold P <5 × 10^−8^). We then defined several candidate variant sets based on additional filtering criteria, including allele frequency and whether insertion-deletion variants (indels) were included. To reduce the redundancy among associated variants, we pruned the correlated signals using chromosome specific LD tag files generated in *PLINK v2.00a3LM* from imputed variants filtered at INFO >0.99 and MAF ≥0.005. LD relationships were defined using r^2^ threshold of 0.8 within 100 kb window in *PLINK v1.90b6.21*. Within each metabolite, variants belonging to the same LD group were compared and the variant with the strongest association signal was retained. Finally, we merged the retained variants across metabolites into a union predictor set.

For the main prediction analyses, we used the set restricted to variants with INFO ≥0.99, MAF ≥0.005, excluding indels and with −log_10_(P) >5. This resulted in a final union set of 13,049 variants for the prediction models.

### Dataset preprocessing for prediction models

For the prediction models, we combined the final set of predictor SNPs with the individual level phenotype and covariate data. We restricted the final phenotype dataset to directly measured, non-derived metabolites to avoid redundancy from derived ratios and composite measures. High-density lipoprotein cholesterol (HDL_C) and low-density lipoprotein cholesterol (LDL_C) were additionally retained because of their clinical relevance, resulting in 109 prediction traits in total. Missingness across metabolites was low, with a maximum of 2.4% (**Table S7**). For each metabolite, missing values were imputed by randomly drawing from the observed values in training data. Missing BMI values (n =77) were handled in the same way. A total of 5,234,004 genotype entries (∼0.43% of all genotype values) were missing prior to imputation. These were imputed separately for each variant using the most common genotype observed in the training set. The genotype values were then coded as −1 (homozygous reference), 0 (heterozygous), and 1(homozygous alternate/effect allele). Before fitting the models, we standardised genotype values, continuous covariates and metabolite traits using statistics from the training set only and then applied the same transformation to the validation and test data. Sex was kept as a binary covariate without scaling.

### Null-label dataset

As a negative control, we constructed a null-label training set by independently permuting each phenotype across individuals within the training set. Genotype data, covariates and all validation and test sets were left unchanged. This preserves the marginal distribution of each phenotype while eliminating any true association between predictors and phenotypes.

### Linear prediction models

We fitted three linear baseline models separately for each phenotype: ordinary least squares^51,52^ (OLS), ridge regression^53,54^ and elastic net regression^55^ predictor set comprised the final set of predictor SNPs together with individual-level covariates. We carried out the analyses both with and without BMI, while age, sex, and the first ten genetic PCs were included throughout. All models were trained using the training set only, with tuning parameters for ridge and elastic net chosen by cross-validation within the training data. Standardisation of genotype values ensured that the regularisation penalty was applied to SNP effects on a comparable scale. While OLS does not require penalisation, the same standardisation was maintained to ensure consistency across all baseline comparisons. Further implementation details, including hyperparameter grids and model configurations, are available in the accompanying code repository (see **Data availability**).

### Residual decomposition of phenotype values for neural network models

For the neural network (NN) analyses, we decomposed metabolite values into sequential components corresponding to covariate effects, genotype effects, and the remaining signal. Let *Y* ∈ ℝ*^nxp^* denote the matrix of metabolite values for *n* individuals and *p* metabolites, *C* the covariate matrix, and *G* the genotype matrix. Conceptually, we considered the additive decomposition

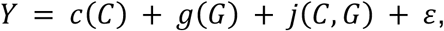

Where *c*, *g* and *j* denote unknown functions mapping their respective inputs to *p* dimensional metabolite predictions and *ε* denotes unexplained residual variation. In practice, these functions were estimated using neural-network models, denoted *ĉ*, *ĝ* and *ĵ*.

We first trained a covariate-only model *ĉ*(*C*) to estimate the component of metabolite variation explained by covariates. For the training set, residuals were constructed using five-fold out-of-fold (OOF) predictions. Specifically, each training individual was predicted by a covariate model trained on the other four folds, yielding 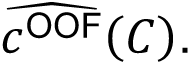 The first-stage residuals were then defined as

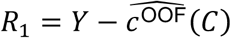

This OOF procedure was used to prevent overly optimistic residual estimates that could arise if each training individual were predicted by a model fitted using that same individual.

We next trained a genotype-only model to predict *R*_1_ from *G*. Again, for the training set, genotype predictions were obtained out-of-fold, yielding 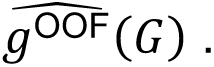 The second-stage residuals were defined as

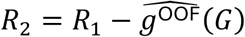

Thus, *R*_2_ represents the component of metabolite variation not captured by covariates or genotype main effects in the first two stages. This remaining signal may include covariate-genotype interactions, other non-additive structure, and residual variation not explained by the preceding models.

In the final stage, we trained a joint model using both covariates and genotypes as input, with *R*_2_ as the target. After the residual targets had been constructed using out-of-fold predictions, final models for each module were fitted on the full training set. For validation and test individuals, predictions were obtained by applying the fitted models sequentially and summing the three components:

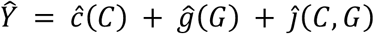

### Neural network training and hyperparameter search

We implemented NN models to capture covariate, genetic, and the remaining effects within the residual decomposition framework described above, considering both multi-task^15^ and single-task settings. In the multi-task NN, a single model was trained to jointly predict all 109 metabolites, with one output node per metabolite. The loss was computed as the mean squared error (MSE) averaged across metabolites. Each module was modelled using a fully connected multilayer perceptron^56^ with ReLU^57^ activations. In the final multi-task NN used for the main analysis, the covariate module had two hidden layers with 125 nodes each, the genotype module had two hidden layers with 512 and 256 nodes, and the joint module had three hidden layers with 1024, 256 and 128 nodes. The input dimensions were 13 covariates for the covariate module and 13,049 genetic variants for the genotype module, while the joint module used concatenated covariates and genetic variants as input.

All models were trained using mini-batch optimisation with AdamW^58^ and early stopping based on validation loss. For the final multi-task NN, the batch size was 64 and the maximum number of epochs was 200 for each module. The learning rates were 0.003, 2 × 10⁻⁵ and 3 × 10⁻⁵ for the covariate, genetic and joint residual modules, respectively. The corresponding weight decay values were 0.1, 0.001 and 0.01, and dropout rates were 0.1, 0.3 and 0.2. Batch normalisation was used only in the covariate module. Training and validation loss curves for the final multi-task NN are shown in **Figure S9**.

To assess the contribution of nonlinear transformations, we used two activation-free multi-task models. First, we constructed a mirror activation-free model by removing nonlinear activation functions from the final multi-task NN while keeping the architecture and hyperparameters unchanged. This provided a direct comparison in which the presence or absence of nonlinear activations was the main difference. Second, we trained an optimised activation-free multi-task model, in which the same stage-wise hyperparameter search procedure (explained below) was applied to the activation-free architecture. This second model was included to avoid comparing the nonlinear NN against an activation-free model that was disadvantaged by hyperparameters selected for the nonlinear setting. In the single-task setting, separate models were trained for each metabolite. For these single-task NNs, we used a configuration selected from a preliminary search on HDL_C (high-density lipoprotein cholesterol). This configuration was applied uniformly across all metabolites. Finally, as a negative control, we trained the same multi-task model on a null-label dataset (**see Methods–Null-label dataset section**), in which phenotype values were independently permuted across individuals within the training set while genotypes, covariates and validation/test data were kept unchanged.

Hyperparameters were optimised sequentially using the model-selection test sets. After a preliminary search, we tuned the covariate module by evaluating combinations of hidden layer dimension, learning rate, weight decay^58^, dropout^59^, and batch normalisation^60^ while keeping the remaining settings fixed. The best-performing configuration was then carried forward to the genotype module, and the same procedure was repeated for the joint module. This stage-wise optimisation was applied to both the multi-task model and the activation-free multi-task architecture, resulting in six module-specific grid searches in total (**Tables S8-10** and **Table S11-13**, respectively).

### SNP-heritability-based reference for genotype-based predictive correlation

To contextualise the magnitude of genotype-based prediction, we compared the observed correlation of the genotype module derived prediction with an approximate reference value derived from external LDSC-based SNP-heritability estimates, following the idea that the variance explained by SNPs constraints the maximum achievable genotype-based prediction accuracy^61^.

Let *y* denote the observed phenotype and the *ŷ_G_* component of the prediction attributable to genotype. The Pearson correlation between prediction and phenotype is

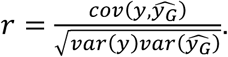

Assuming that the predictor captures only the genetic component (*G*) of the phenotype and that estimation noise (*E*) is negligible, the variance of the prediction approximates the genetic variance, *var*(*ŷ_G_*) ≈ *var*(*G*). Assuming *cov*(*G*, *E*) = 0, the covariance between the phenotype and the genetic component is

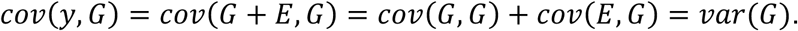

In this setting, the expected maximum correlation is

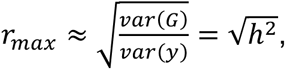

where ℎ^2^ denotes SNP-heritability on the observed scale.

We obtained metabolite-specific SNP-heritability estimates from a large-scale GWAS of circulating metabolites^39^. Because these estimates were derived using LD score regression and reflect genome-wide additive SNP effects rather than the variance explained by the selected predictor variants here, we interpreted 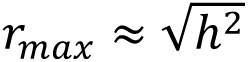 as an approximate reference ceiling under additive model rather than a strict bound. For each metabolite, we compared this expected value with the Pearson correlation between the genotype-module prediction and the observed phenotype in the held-out test set.

### Model evaluation

Model performance was evaluated on the held-out test set. For each metabolite, we calculated the coefficient of determination (*R*^2^), mean squared error (MSE), mean absolute error (MAE), and Pearson correlation coefficient (r) between observed and predicted metabolite values. *R*^2^ was calculated as 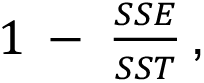 where *SSE* is the residual sum of squares between observed and predicted values and *SST* is the total sum of squares around the observed test-set mean. MSE and MAE were calculated as the mean squared and absolute prediction errors, respectively. All metrics were computed separately for each metabolite and then summarised across metabolites.

For the residual NN models, we additionally evaluated module-level performance. First, we calculated the predictive performance of the covariate module alone. We then recalculated performance after adding the genotype module prediction to the covariate-module prediction. The difference between these two values was used to estimate the incremental gain from the genotype module. Finally, we recalculated performance after adding the joint residual module to the covariate and genotype module predictions; the difference between this full-model performance and the covariate-plus-genotype performance was used to estimate the incremental gain from the joint residual module.

In addition to these incremental gains, we also evaluated the isolated genotype and joint module predictions by comparing each module’s prediction directly with the observed metabolite values. These isolated module metrics were used only to assess whether the residual components were aligned with the phenotype. They should not be interpreted as incremental variance explained, because the genotype and joint modules were trained on residual targets rather than directly on the original metabolite values.

### Bootstrap-based uncertainty estimation

Uncertainty in performance metrics was estimated using bootstrap resampling at the individual level. For each replicate, individuals from the held-out test set were sampled with replacement, and all metrics were recomputed on the same resampled individuals. For model comparisons, the bootstrap was paired: the same resampled individuals were used for each model within a replicate, and performance differences were calculated within that replicate. This procedure was applied consistently across analyses, including overall performance summaries, component-wise evaluations, and model comparisons. We performed 1,000 bootstrap replicates and report 95% confidence intervals based on empirical percentiles of the bootstrap distribution.

## Funding

M.N.G, B.Y. were supported by the Estonian Research Council grant PSG1179. M.N.G., L.M. were supported by the Estonian Research Council grant PRG2625. M.A. was supported by the Estonian Research Council grant PSG1028. T.H. was supported by the Estonian Research Council grant PRG2585. F.J. was supported by ANR-20-CE45-0010 RoDAPoG. The research was conducted using the Estonian Center of Genomics/Roadmap II funded by the Estonian Research Council (project number TT17). The research was conducted using the research infrastructure ‘Estonian Center for Genomics’ funded by the Estonian Research Council (project number TARISTU24-TK19).

## Acknowledgements

We thank members of the Research group of pharmacogenomics for helpful discussions and feedback throughout the project. We thank Ralf Tambets and Kaur Alasoo for providing the SNP-heritability estimates and Laura Birgit Luitva for providing the GEVA snapshot data used in this study. Data analysis was carried out in part in the High-Performance Computing Center of University of Tartu. We acknowledge the Estonian Biobank Research Team for providing the genotype, phenotype and metabolomics data resources used in this work. In particular, we thank Andres Metspalu, Lili Milani, Tõnu Esko, Reedik Mägi, Mait Metspalu, Mari Nelis and Georgi Hudjashov for their contributions to data collection, genotyping, quality control and imputation, and Priit Palta, Nele Taba, Erik Abner, Jaanika Kronberg and Urmo Võsa for their contributions to the generation, development and quality control of the Nightingale metabolomics data layer.

## Author Contributions

Conceptualisation: M.N.G., B.Y. Methodology: M.N.G., B.Y. Software: M.N.G. Formal Analysis: M.N.G., B.Y. Investigation: M.N.G. Data Curation: M.N.G., Estonian Biobank Research Team. Original Draft: M.N.G. Review & Editing: M.N.G., M.A., T.H., F.J., L.P., L.M., B.Y. Visualisation: M.N.G. Supervision: F.J., L.P., L.M., B.Y. Project Administration: B.Y. Funding Acquisition: L.M., B.Y. All authors reviewed and approved the final manuscript.

## Materials & Correspondence

Correspondence and requests for materials should be addressed to Burak Yelmen or Merve Nur Güler.

## Data availability

Code for this study is available at https://github.com/genodeco/Multi-task-mets.

## Competing interests

The authors declare no competing interests.

## Ethics statement

The activities of the EstBB are regulated by the Human Genes Research Act, which was adopted in 2000 specifically for the operations of the EstBB. Individual level data analysis in the EstBB was carried out under ethical approval 1.1-12/624 (issued 24 March 2020), 1.1-12/2618 (issued 04 August 2022), and 1.1-12/3454 (issued 20 October 2022) from the Estonian Committee on Bioethics and Human Research (Estonian Ministry of Social Affairs).

## Notes

### Competing Interest Statement

The authors have declared no competing interest.

### Summary of Updates

Abstract and end of the Introduction section have been updated.

