## Supplemental Figures for "Covariate-aware genomic prediction of blood metabolite profiles using multi-task neural networks"

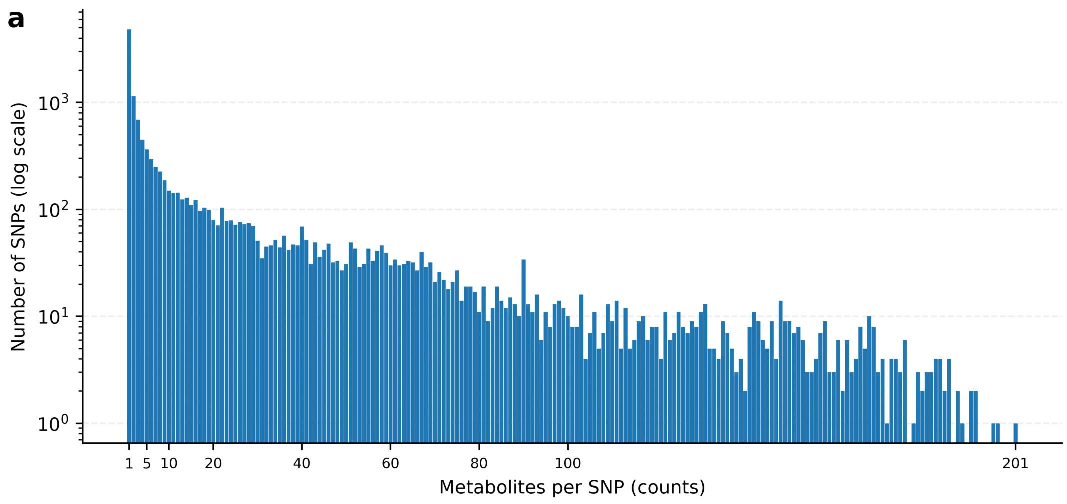


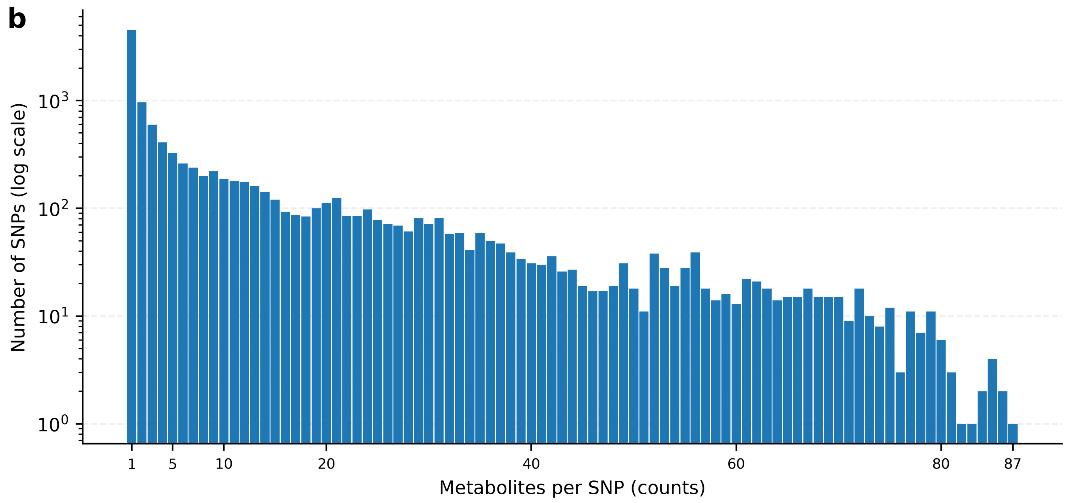


**Figure S1. Distribution of cross-metabolite sharing in the genetic predictor set. a)** Histogram showing the number of metabolite associations per SNP across the full set of 249 Nightingale metabolomics measures for the 13,049 variants included in the final predictor set. The predictor set was defined as the union of variants $-\log_{10}(P) > 5$ (=P $<1\text{x}10^{-5}$) across metabolite-specific GWAS performed in the training set, followed by LD pruning and merging across metabolites. Association counts were calculated across all 249 metabolites because the final predictor set was constructed from the full metabolomics panel, including derived biomarkers, rather than only from the 109 prediction targets. The y-axis is shown on a logarithmic scale. **b)** Same distribution after restricting the association counts to the 109 metabolites used as prediction targets. This restricted view is shown for comparison, but it excludes approximately 2,000 variants that entered the final predictor set through associations with derived metabolites. The y-axis is shown on a logarithmic scale.

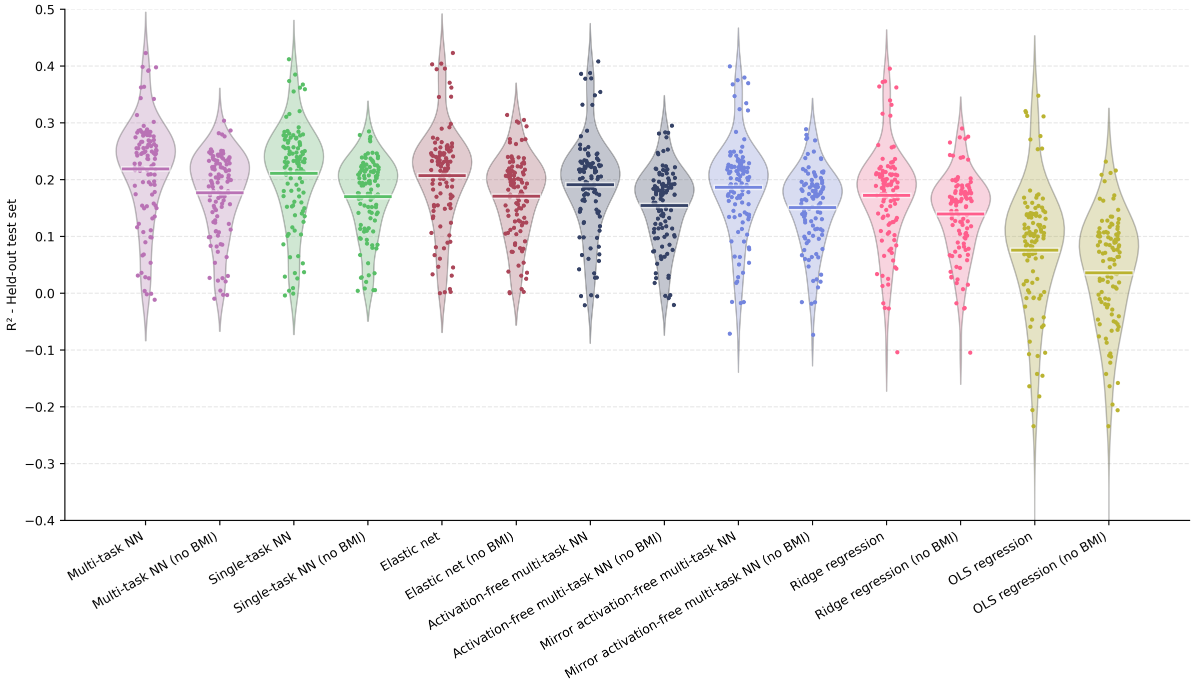


**Figure S2. Comparison of model performance with and without BMI as an input covariate.** Violin plots show the distribution of held-out test-set $R^2$ values across metabolites for each model fitted with BMI included or excluded from the covariate set. Points represent individual metabolites and horizontal bars indicate model means. The y-axis is truncated to improve visibility of the main distribution; one extreme negative OLS value falls below the plotted range.

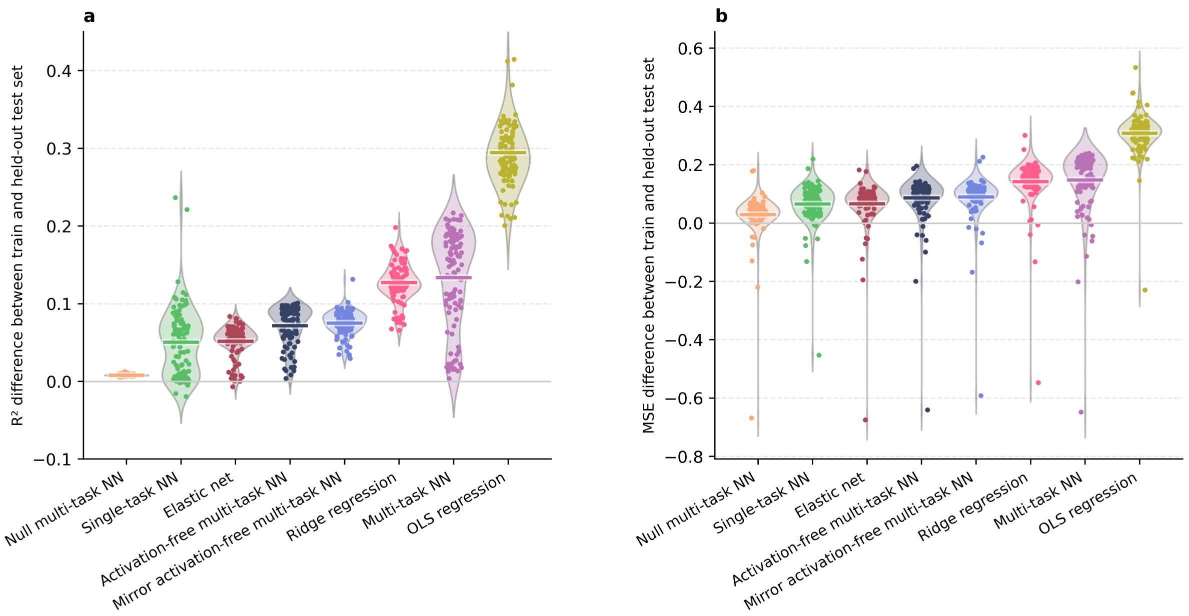


**Figure S3. Difference between training and held-out test performance across models. a)** Distribution of the per-metabolite difference in $R^2$, defined as training $R^2$ minus held-out test $R^2$. Positive values indicate higher performance on the training set than on the held-out test set. **b)** Distribution of the per-metabolite difference in mean squared error (MSE), defined as held-out test MSE minus training MSE. Positive values indicate larger prediction error on the held-out test set. MSE was calculated on the training-standardised phenotype scale; however unlike $R^2$, it is not divided by the phenotype variance within each evaluation split. Large MSE gaps can therefore partly reflect split-specific differences in the phenotype variance, especially for skewed metabolites with extreme tail values (**Table S14**).

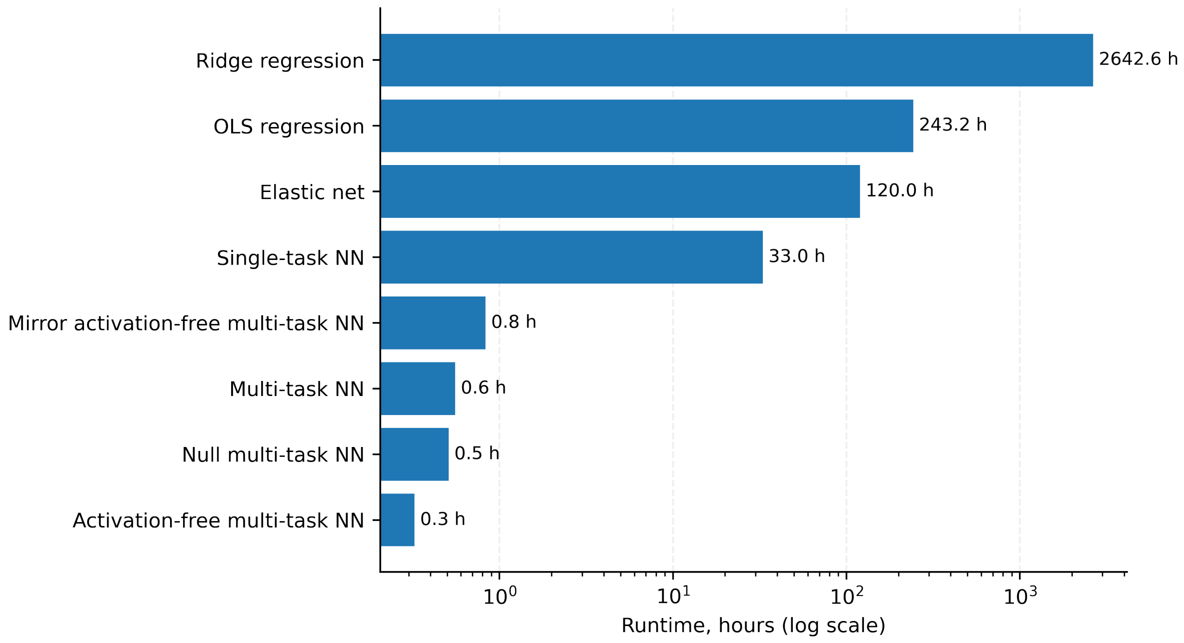


**Figure S4: Runtime comparison across model classes.** Runtime was summarised as the cumulative wall-clock time. Multi-task neural networks were trained as single multi-output jobs, whereas the single-task neural network was trained separately for each metabolite; its runtime therefore represents cumulative compute time across 109 jobs rather than elapsed calendar time. Linear models were run on the CPU partition, while neural-network models were run on one Tesla V100 GPU.

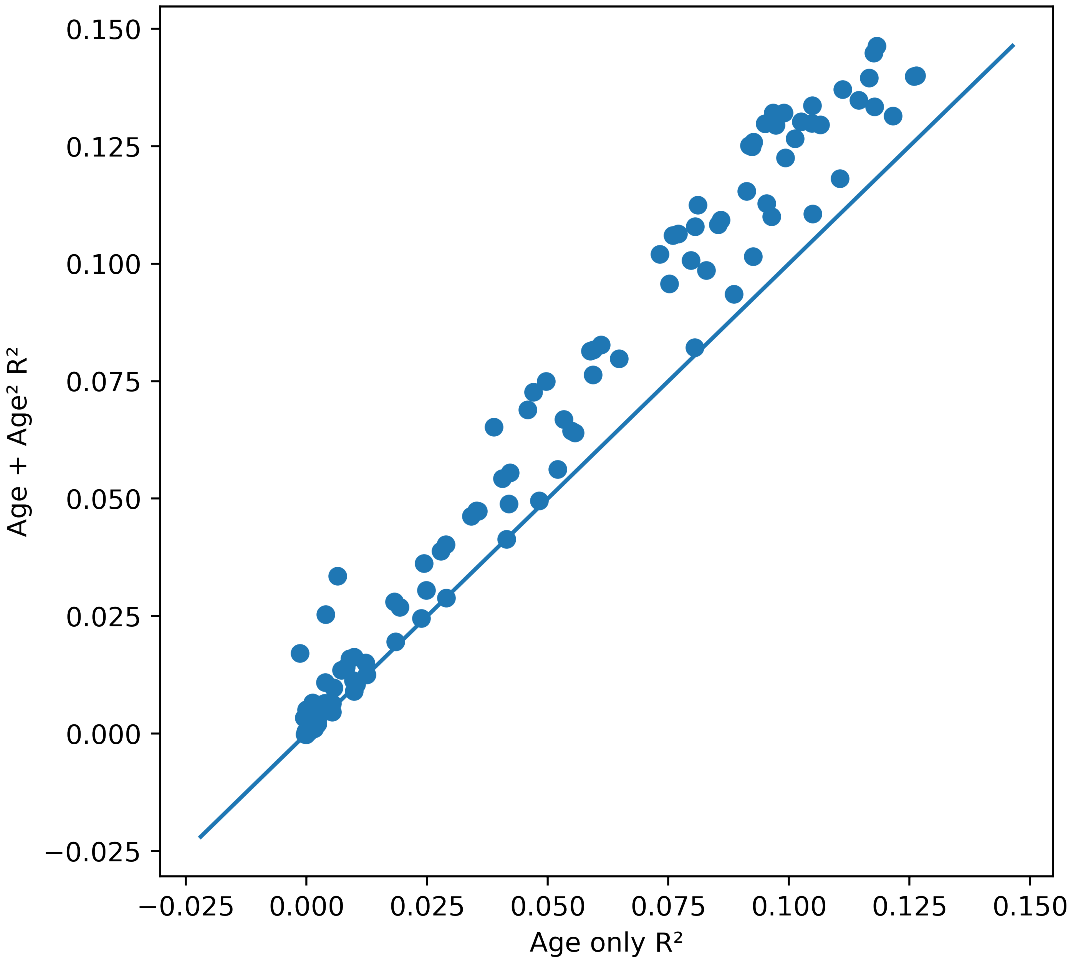


**Figure S5. Improvement in covariate modelling with a quadratic age term.** Each point represents one metabolite. The x-axis shows held-out test-set $R^2$ from a linear model including age only, and the y-axis shows $R^2$ from a model including both age and age$^2$. The diagonal indicates equal performance. Most metabolites lie above the diagonal, indicating consistent improvement when allowing simple nonlinear covariate effects.

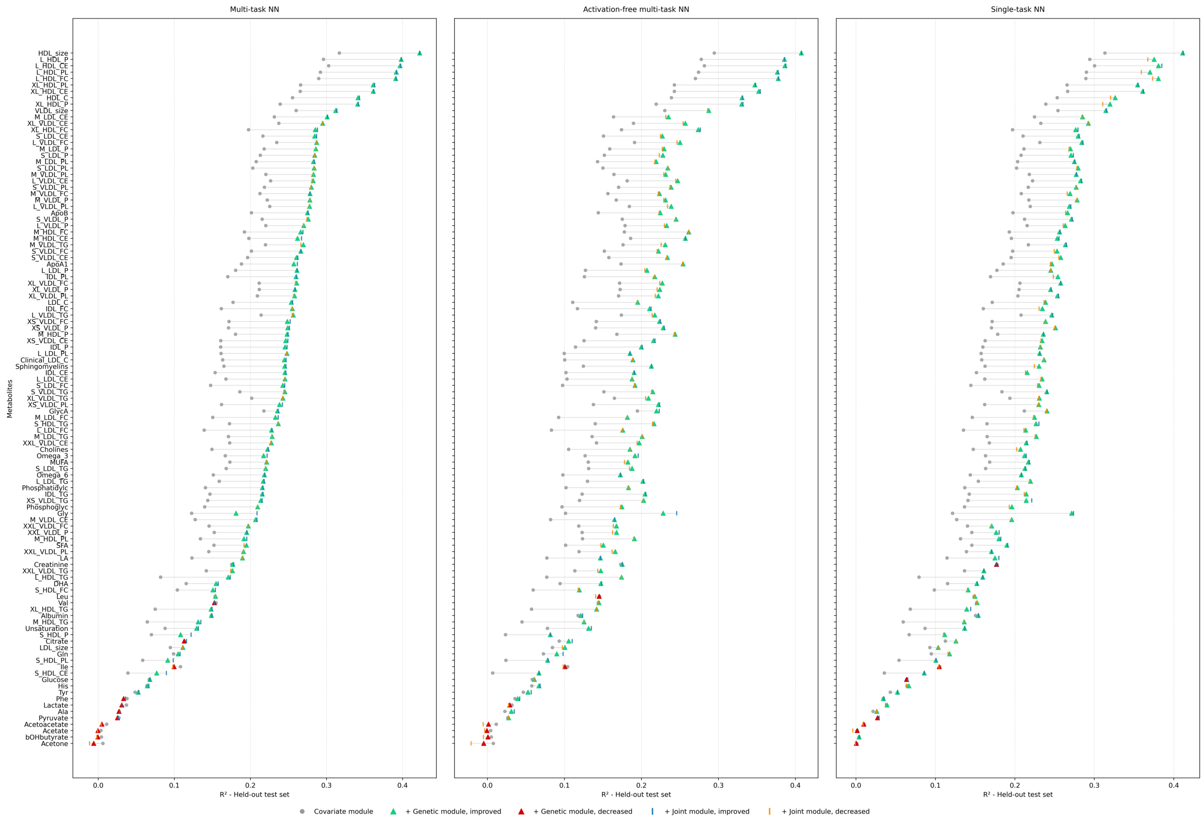


**Figure S6. Metabolite-level accumulation of predictive performance across decomposition stages.** For each metabolite, grey circles indicate held-out test-set $R^2$ for the covariate module alone. Triangles show performance after addition of the genetic module, with green triangles indicating improvement relative to the covariate module and red triangles indicating a decrease. Vertical marks show performance after inclusion of the final joint residual module, with blue marks indicating improvement relative to the covariate-plus-genetic stage and orange marks indicating a decrease. Metabolites are ordered by full-model performance based on multi-task neural network.

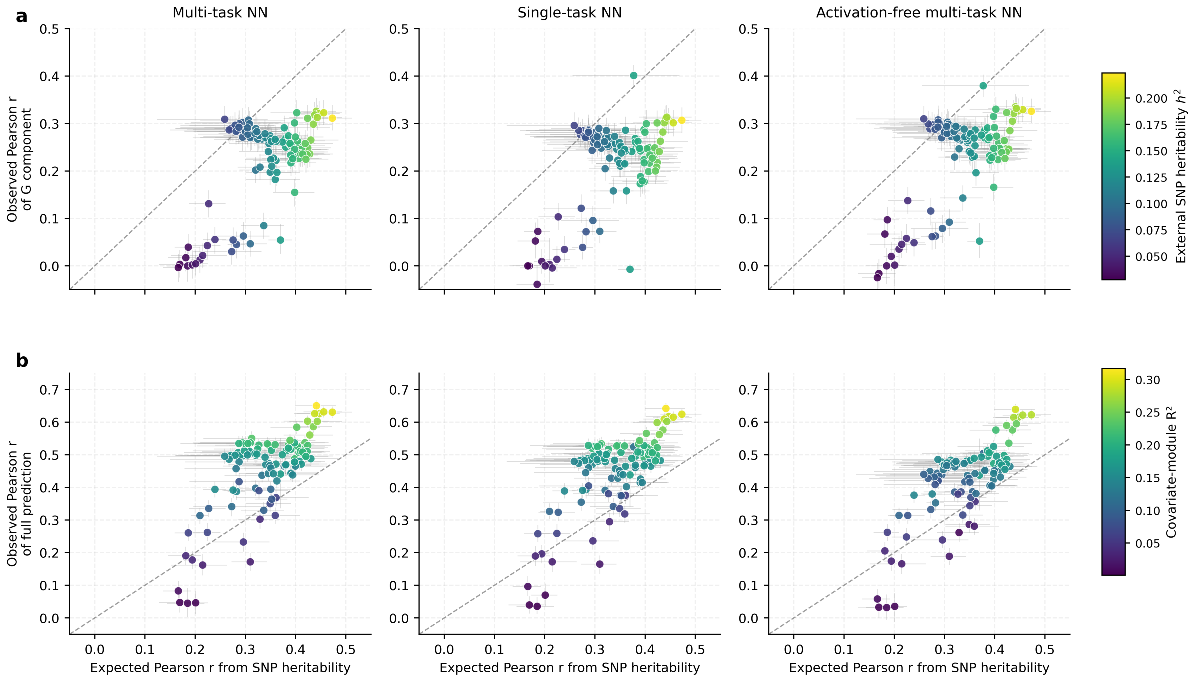


**Figure S7: SNP-heritability-derived genetic expectation compared with genotype component and full-model predictive correlation across metabolites**. Each point represents one metabolite. The x-axis shows the expected maximum Pearson correlation derived from external LDSC-based SNP-heritability estimates, calculated as $\sqrt{h^2}$, and used as an approximate genetic reference. The y-axis shows the observed Pearson correlation in the held-out test set **a)** for the genotype-derived prediction component alone and **b)** for the full model prediction. Columns show results for the multi-task neural network, single-task neural network and activation-free multi-task neural network. Horizontal intervals indicate uncertainty in the expected correlation propagated from the standard error of the external LDSC-based SNP-heritability estimate, whereas vertical bars indicate 95% confidence intervals obtained from paired bootstrap resampling of individuals for the observed correlation. In panel **a**, points are coloured by the external SNP-heritability estimate. In panel **b**, points are coloured by covariate-module $R^2$. The dashed diagonal line indicates equality between the SNP-heritability-derived genetic expectation and the observed predictive correlation.

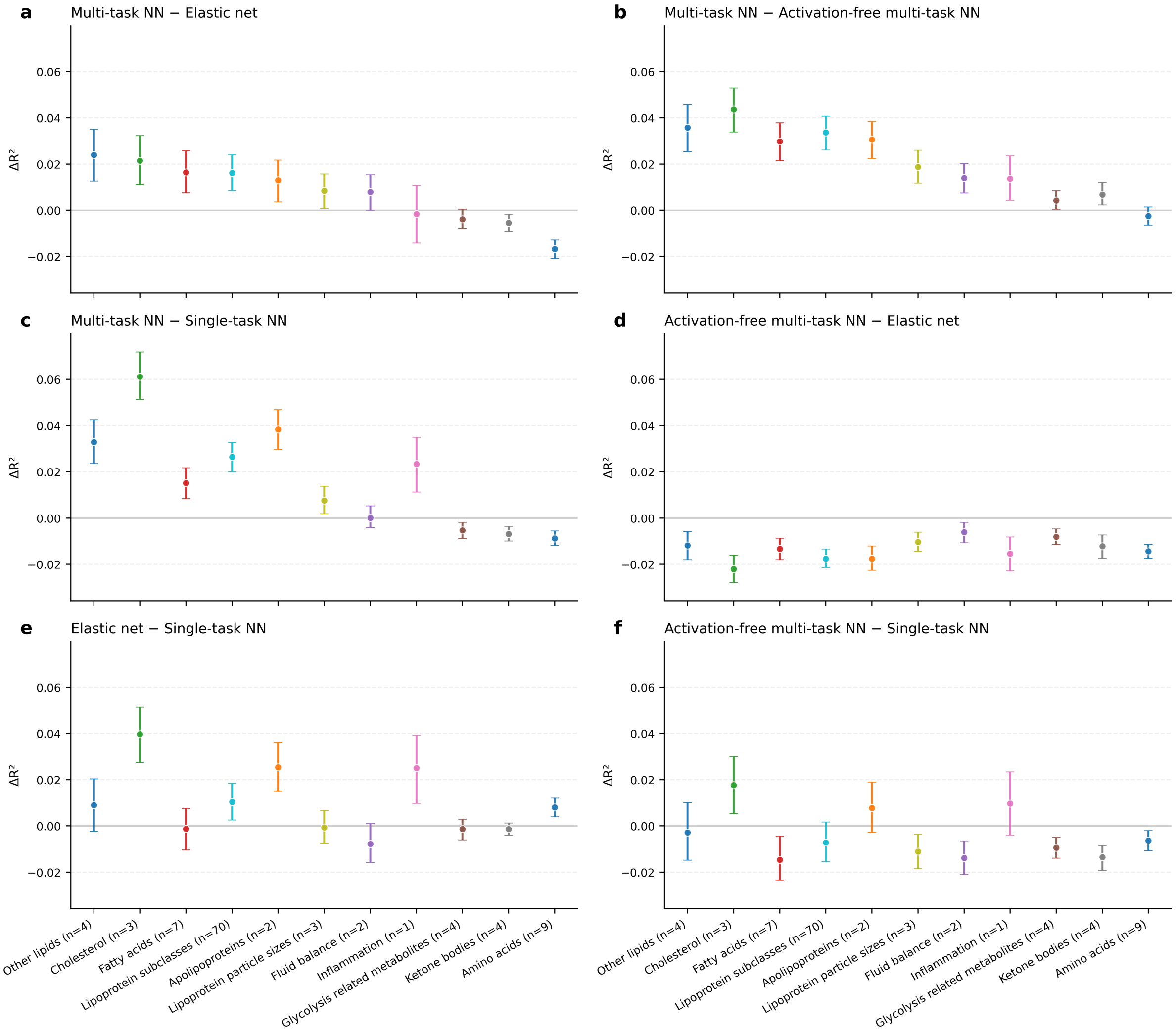


**Figure S8. Group-wise difference in held-out $R^2$ between the multi-task neural network and comparator models.** Panels show the group-level mean difference in held-out test-set $R^2$ for **a)** multi-task neural network (NN) - elastic net, **b)** multi-task NN - activation-free multi-task NN, **c)** multi-task NN - single-task NN, **d)** activation-free multi-task NN - elastic net, **e)** elastic net - single-task NN and **f)** activation-free multi-task NN - single-task NN. Points indicate the group-wise mean $\Delta R^2$ and error bars indicate 95% confidence intervals estimated by paired bootstrap resampling over individuals (n =1,000). Positive values indicate better performance of the first model named in each panel title. Labels on the x-axis indicate the number of metabolite traits in each group.

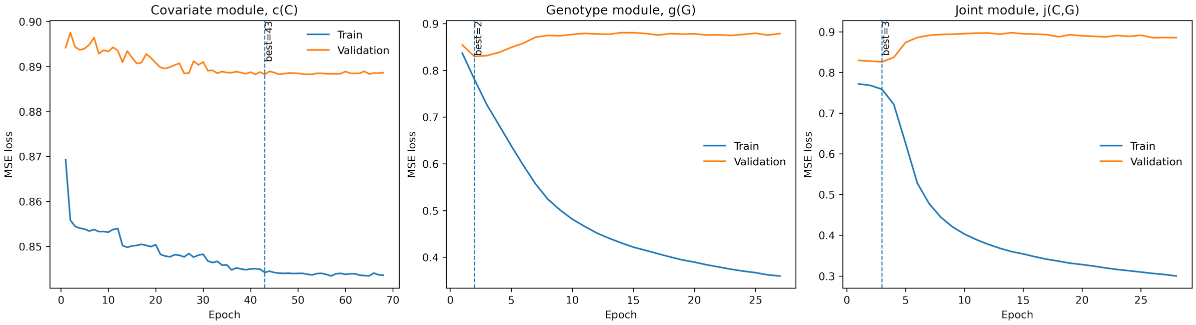


**Figure S9: Training and validation loss curves for the multi-task neural network.** Loss curves are shown for the three modules of the residual decomposition framework: covariate module, genotype module and joint module. Blue lines show training loss and orange lines show validation loss across epochs. Dashed vertical lines indicate the epoch selected by early stopping based on validation loss. The covariate module showed a gradual reduction in validation loss, whereas the genotype and joint residual modules reached their best validation loss early despite continued decreases in training loss.
